# Women’s Decision-Making Autonomy and ICT Utilization on Access to Antenatal Care Services: Survey Results From Jinja and Kampala Cities, Uganda

**DOI:** 10.1101/557413

**Authors:** Hasifah K. Namatovu, Tonny J. Oyana, Jude T. Lubega

## Abstract

There is growing evidence in Uganda that the non-attendance of antenatal care is largely influenced by the lack of decision-making autonomy, inadequate information and poor services offered in health facilities. Although previous studies have examined barriers and facilitators of antenatal care, a few of them have investigated the extent of decision making autonomy and ICT adoption among expectant mothers. A cross sectional design through focus group discussions and survey questionnaires was used to collect data. Three hundred and twenty households were randomly sampled in Kampala and Jinja districts. The Chi-square tests (χ^2^) for independence to analyze group differences among women’s socio-demographic characteristics and decision-making autonomy was used. Inclusion criteria included respondents aged 18 and 50 years, completion of primary school education, expectant mothers and mothers who gave birth two years prior to the study. A hundred and sixty-four respondents participated in this survey. About 59.5% of women lacked decision making autonomy. Midwives (37.6%) and village health teams (35%) were a major source of antenatal care information, and 49.5% of expectant mothers lacked ANC information. Ninety percent (90%) of mothers did not use any form of ICT’s to enhance their decisions yet 79% possessed mobile phones. We observed a strong association between antenatal care decision-making autonomy and women with higher education (χ^2^ = 8.63, *ρ* = 0.035), married (χ^2^ = 4.1, *ρ* = 0.043) and mature (36–50) (χ^2^ = 8.81, *ρ* = 0.032). The main findings in this study suggest that ICT adoption and decision making autonomy among expectant mothers is still low and less appreciated. Control measures and interventions should be geared towards empowering women to influence their decisions.

## Background

Maternal mortality is still high and remains a big public health issue among countries situated in Sub-Saharan Africa [1, 2]. It is estimated that 358,000 women (830 deaths everyday) died worldwide in 2008, as a result of preventable causes related to pregnancy and childbirth [3]. This is more than 34 women per hour. Of these, 99% occurred in developing countries. Regionally, Sub-Saharan and South Asia accounted for 87% of the global maternal deaths [4]. In Uganda an estimated 16 women die from giving birth every day; on average this is one death every hour and a half, and nearly 6,000 every year [5, 6]. High maternal mortality is attributed to lack of access to modern family planning, low awareness of danger signs of pregnancy/labour [2, 7]. Furthermore, the inability of some pregnant women to attend antenatal care (ANC), deliver in a health facility and facility location has aggravated the situation [7, 8].

Many of these deaths are preventable once mothers get adequate maternal access and emergency obstetric care (EmOC) [9]. The most critical intervention of maternal mortality is, i) participating in ANC, ii) delivery by skilled birth attendant (SBA), iii) access to EmOC, and iv) access to family planning services [10, 11].

ANC establishes the first contact with health facilities and it is highly premised that mothers who have attended at least more than one ANC are more likely to give birth with a help of a SBA [12]. ANC provides an avenue for mothers to receive information and guidance on safe childbirth and help prevent, detect and alleviate health problems that affect mothers and babies during pregnancy [7, 13, 14]. WHO recommends every pregnant woman to visit a health facility at least four times during pregnancy [15]. Despite numerous efforts to have women attend ANC in Uganda, only 47% attended the recommended four visits while 58% delivered in a health facility [16, 17]. However, little information and knowledge is available on the barriers and facilitators of ANC in rapidly developing urban centers. The purpose of this study is to investigate the extent of ICT adoption and decision making autonomy in influencing access to ANC in Uganda.

## Methods and Materials

### Study Setting and Sampling

Kampala and Jinja population represent a typical urban population that is rapidly growing with high demand for high-quality ANC services, and provides a representative sample to understand urban health among women. A mixed method (quantitative and qualitative) approach was used to collect and analyse the evidence received from respondents. Through purposive and stratified random sampling techniques, this study ensured that expectant mothers and mothers who gave birth two years prior to the study were properly drawn from sparsely and densely populated neighbours in Kampala and Jinja districts. The use of these two sampling techniques yielded a representative subset of all women receiving ANC. The sample was used to ascertain the type and nature of decisions that mothers routinely engage in while pregnant. Inclusion criteria included respondents aged between 18 and 50 years old, completion of primary school education, expectant mothers and mothers who gave birth two years prior to the study.

### Study Design

A cross sectional design was used to collect baseline and thematic data on the study population using a two-step approach. The sample population was drawn from Walukuba, Mpumudde, Mengo Kisenyi and Kasubi Kawaala parishes situated in Jinja and Kampala district covering a period from January to April 2015. The first step involved the creation of baseline data through interviewing 35 expectant mothers from 59 randomly sampled households. Baseline data was used to understand the characteristics of the sample population and this informed an in-depth study. In Walukuba, a total of 8 mothers were interviewed, N=3 were pregnant and N=5 were mothers with children below two years. In Mpumudde, 6 women were interviewed, N=2 were pregnant and N=4 were mothers with children below two years. In Mengo Kisenyi, 10 participated, N=2 were pregnant and N=8 were mothers with children below two years. In Kasubi Kawaala, 11 participated, N=3 were pregnant and N=8 were mothers with children below two years. Baseline studies are important for establishing key priority areas of the study and provide quantitative information on the current population status [18].

The second step involved the collection of thematic data through a survey questionnaire. Using baseline data, a survey questionnaire was formulated based on three major themes (demographics; source and type of information that aids decision making, decision making autonomy and barriers of access to ANC services; and uptake of ICT). A survey questionnaire with open-ended and closed-ended questions was pre-tested on women residing in Lubya zone, which is outside the surveyed area.

A stratified random sample of 320 households located in different strata in Jinja and Kampala was used to administer 350 survey questionnaires. The sampling took place in three phases; defining the characteristics of study population and partitioning it into households, choosing the sample size based on different strata (a 20 m to 30 m distance between two households, distance to nearest ANC service point and female demographics); and randomly selecting the first household to administer the survey questionnaire.

Two trained research assistants together with one female local resident guided the house to house distribution of questionnaires. Before the survey, they were trained on the primary objective of this study, ethical code of conduct, temperament, especially where respondents were rude and non-responsive. Each sampled household was approached by a research assistant to seek consent from the household head and determine whether there was a pregnant woman or a mother with a child below two years old present. Households that were near the main road or a trading centre were used as a starting point, and subsequent households were selected from the sampling list. This whole process took an average of 48 days.

Survey questionnaires were collected after one week, and women who could not comprehend the content of the questionnaire were guided by the interviewers in a verbal one-on-one interaction. In some cases, these had to be translated in local languages. However, use of ANC cards to capture the number of times a woman had attended ANC, parity and their age was minimally used because most of them, especially those that had given birth prior the survey had either lost or discarded the cards.

In the qualitative study, 10 pregnant women, 5 from each district were carefully selected to participate in the focus group discussion that lasted approximately 80 minutes.

### Data analysis

All data was coded, processed and analyzed using IBM SPSS Statistics Version 25 (New York, USA). Analysis to test the association of socio-demographic and decision-making autonomy was achieved using Chi-square tests (χ^2^).

Qualitative responses were analyzed using conventional content analysis, which involved reading data repeatedly to try and identify common patterns. Any patterns identified from the written statements, oral or audio data were encoded.

#### Ethical Approval

Written consent to conduct this survey was sought from Kampala City Council Authority. Further approval was orally sought from the local council leaders. All women who participated gave a verbal consent while others sought consent from their spouses.

## Results

### Descriptive results

Socio-demographics of the study population is summarised in the Table 1. The total of 320 households were visited, 145(45.4%) in Jinja and the rest in Kampala. Thirty three percent of the respondents had completed primary school education, 36.2% secondary, 11.7% vocational while 19% had done university. At the time of the study, 42(25.6%) women were pregnant while 122(74.4%) were mothers who had given birth two years prior the study. Two fifth (39.5%) of the respondents had given birth to two children, 40(24.4%) had three, 26(15.9%) had one, 21(12.8%) had more than four while 12(7.3%) had no child but were pregnant. Of the households visited, three quarters (75.9%) were headed by men; 24.1% by women; 151(47.3%) had a 5km or more distance from a public health facility while 169(52.7%) from a private health facility.

**Table 1:** Demographics

| Number that participated and returned questionnaires | Number/Percent of Mothers<br>N=164 |  | Total |
| --- | --- | --- | --- |
|  | Kampala | Jinja |  |
| Number of Respondents | 83 | 81 | 164 |
| Households visited | 175(54.6%) | 145(45.4%) | 320(100%) |
| Education Level |  |  |  |
| Primary | 19(11.7%) | 35(21.5%) | 54(33.1%) |
| Secondary | 28(17.2%) | 31(19%) | 59(36.2%) |
| Vocational | 17(10.4%) | 2(1.2%) | 19(11.7%) |
| University | 18(11%) | 13(8%) | 31(19%) |
| Age Group |  |  |  |
| 18–25 years | 30(18.3%) | 52(31.7%) | 82(50%) |
| 26–35 years | 38(23.2%) | 20(12.2%) | 58(35.4%) |
| 36–45 years | 12(7.3%) | 7(4.3%) | 19(11.6%) |
| 46–50 years | 3(1.8%) | 2(1.2%) | 5(3%) |
| Status |  |  |  |
| Pregnant | 23(14%) | 19(11.6%) | 42(25.6%) |
| Mother with baby less than two years | 60(36.6%) | 62(37.8%) | 122(74.4%) |
| Parity |  |  |  |
| None | 7(4.3%) | 5(3%) | 12(7.3%) |
| One | 14(8.5%) | 12(7.3%) | 26(15.9%) |
| Two | 27(16.5%) | 38(23.2%) | 65(39.5%) |
| Three | 22(13.4%) | 18(11%) | 40(24.4%) |
| Four and above | 13(7.9%) | 8(4.9%) | 21(12.8%) |
| Marital Status |  |  |  |
| Married or co-habiting | 63(38.4%) | 69(42%) | 132(80.4%) |
| Staying Single | 20(12.4%) | 12(7.2%) | 32(19.6%) |
| Working Status |  |  |  |
| Working with income | 29(17.6%) | 38(23.1%) | 67(40.7%) |
| Not working | 54(33%) | 43(26.3%) | 97(59.3%) |
| Household Heads |  |  |  |
| Male | 138(43.2%) | 105(32.7%) | 243(75.9%) |
| Female | 37(11.6) | 40(12.5%) | 77(24.1%) |
| Households within 5km distance or more from a health facility |  |  |  |
| Nearest public health facility | 78(24.5%) | 73(22.8%) | 151(47.3%) |
| Nearest private health facility | 97(30.2%) | 72(22.5%) | 169(52.7%) |

It is also indicated that half of these respondents in this age bracket had given birth to at least one child, many of these (35.3%) having two or more children (See table 2). Further analysis indicates that out of those that had completed primary and secondary education, 25.7% and 28.7% had given birth to two or more children respectively.

**Table 2:**
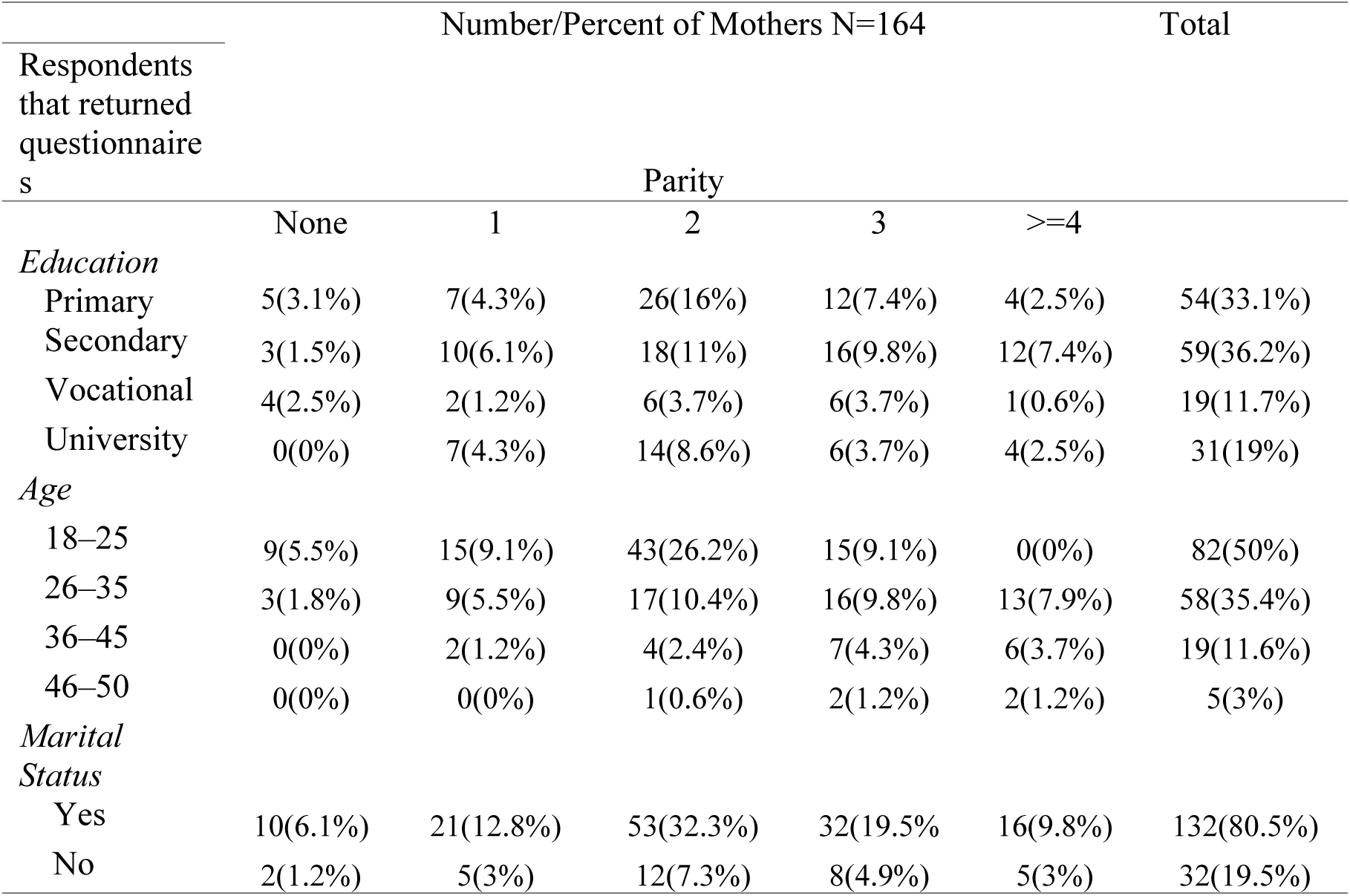
Reproductive health information of expectant mothers

|  | Number/Percent of Mothers N=164 |  |  |  |  | Total |
| --- | --- | --- | --- | --- | --- | --- |
| Respondents<br>that returned<br>questionnaire<br>s | Parity |  |  |  |  |  |
|  | None | 1 | 2 | 3 | >=4 |  |
| *Education* |  |  |  |  |  |  |
| Primary | 5(3.1%) | 7(4.3%) | 26(16%) | 12(7.4%) | 4(2.5%) | 54(33.1%) |
| Secondary | 3(1.5%) | 10(6.1%) | 18(11%) | 16(9.8%) | 12(7.4%) | 59(36.2%) |
| Vocational | 4(2.5%) | 2(1.2%) | 6(3.7%) | 6(3.7%) | 1(0.6%) | 19(11.7%) |
| University | 0(0%) | 7(4.3%) | 14(8.6%) | 6(3.7%) | 4(2.5%) | 31(19%) |
| *Age* |  |  |  |  |  |  |
| 18–25 | 9(5.5%) | 15(9.1%) | 43(26.2%) | 15(9.1%) | 0(0%) | 82(50%) |
| 26–35 | 3(1.8%) | 9(5.5%) | 17(10.4%) | 16(9.8%) | 13(7.9%) | 58(35.4%) |
| 36–45 | 0(0%) | 2(1.2%) | 4(2.4%) | 7(4.3%) | 6(3.7%) | 19(11.6%) |
| 46–50 | 0(0%) | 0(0%) | 1(0.6%) | 2(1.2%) | 2(1.2%) | 5(3%) |
| *Marital*<br>*Status* |  |  |  |  |  |  |
| Yes | 10(6.1%) | 21(12.8%) | 53(32.3%) | 32(19.5%) | 16(9.8%) | 132(80.5%) |
| No | 2(1.2%) | 5(3%) | 12(7.3%) | 8(4.9%) | 5(3%) | 32(19.5%) |

Results in table 3 indicate that more than three quarters (80.5%) of the respondents were married, majority of these ranging between 18–25 years.

**Table 3:**
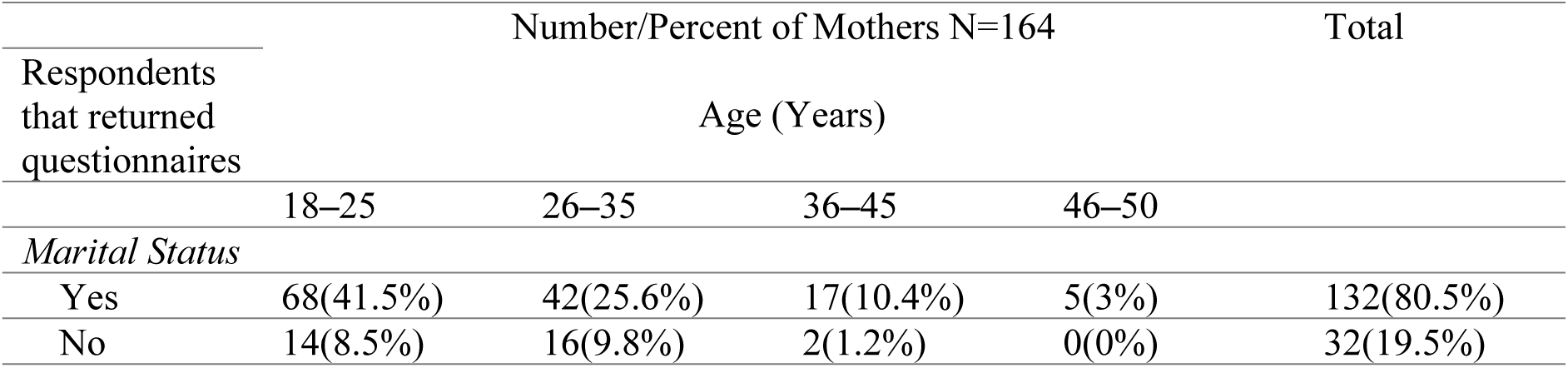
Marital Status and age

The Pearson’s Chi-square results presented in table 4 indicated that there was a significant association between age and parity (χ^2^ = 38.15, *ρ*<0.001), age and education (χ^2^ = 35.12, *ρ*<0.001), decision making autonomy and education (χ^2^ = 8.63, *ρ* = 0.035), decision making autonomy and age (χ^2^ = 8.81, *ρ* = 0.032), decision making autonomy and marital status (χ^2^ = 4.1, *ρ* = 0.043). However, analysis shows that there was no association found between education and parity (χ^2^ = 17.8, *ρ* = 0.122), age and marital status (χ^2^ = 4.9, *ρ* = 0.179), education and marital status (χ^2^ = 3.14, *ρ* = 0.371), decision making autonomy and parity (χ^2^ = 4.29, *ρ* = 0.368).

**Table 4:**
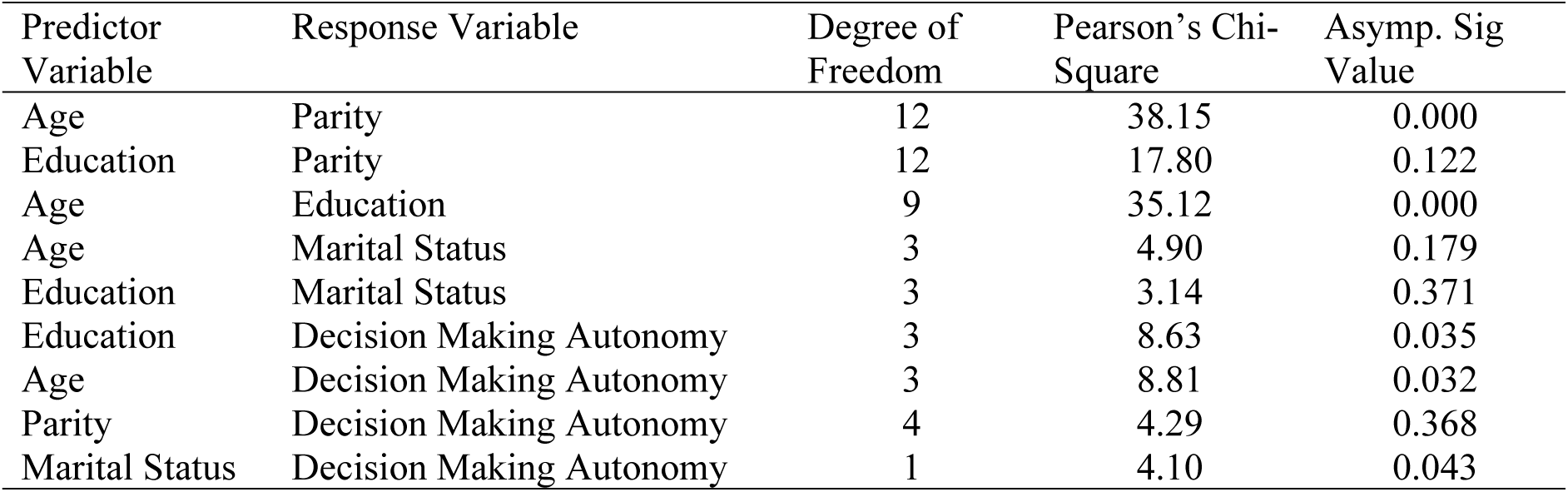
Test of Independence between the Predictor and Response Variable

### Antenatal Care and Decision-Making Practices

#### Major source of antenatal care information

Thirty seven percent of women relied on midwives for ANC information although more than half (24.8%) of these were from Kampala. Slightly less than one third (35%) depended on village health teams (VHT) majority (22.9%) were from Jinja. Nineteen percent depended on peer mothers, 4% depended on family and friends while 3% depended on none other than themselves.

#### Decision-making autonomy

Fifty nine percent of the mothers did not have the autonomy to influence their ANC decision making with majority (34.4%) coming from Jinja.

#### Why mothers did not make their antenatal care decisions?

Lack of knowledge about ANC (49.5%), husbands being the sole financial providers (32%), others did not know (12.4%), culture (1%) and lack of self-trust (1%).

#### Use and Uptake of ICT

Prior to the study, seventy nine percent of the mothers (see figure 2) owned mobile phones although more than three quarters (90%) had never used any ICT’s for ANC (see table 5). Almost ninety percent (89.9%) of the mothers were not using any known ICT’s for their ANC practices. However, a few of those that used ICT’s in ANC, 8% used mobile phones while 1.9% used computers (table 5). Some of the ICT services that were used to access ANC were SMS (2.4%), instant messages (4.3%), voices services (2.4%) (Figure 3). Findings shown in figure 4 indicate that apart from voice and SMS, mother’s used mobile phones for WhatsApp (56.7%), Facebook (36.6%), mobile money (5.5%), twitter (0.6%) and others like Instagram (0.6%). In figure 5, mothers indicated several barriers that limited the uptake of ICT services in supporting their decisions and these included, the high cost of data and voice services (15.2%), poor quality internet services (5.5%), lack of skills to use ICT tools and services (42.1%), limited knowledge on the importance of ICT in ANC (35.4%) and the lack of integrated services (1.8%).

**Table 5:** Antenatal care and decision-making practices of expectant mothers

| Number that participated and returned questionnaires | Number/Percent of Mothers<br>N=164 |  | Total |
| --- | --- | --- | --- |
|  | Kampala | Jinja |  |
| *Major source of ANC information* |  |  |  |
| Peer Mothers | 9(5.7%) | 22(14%) | 31(19.7%) |
| Midwives | 39(24.8%) | 20(12.7%) | 59(37.6%) |
| Family and Friends | 6(3.8%) | 1(0.6%) | 7(4.5%) |
| Village health teams | 19(12.1%) | 36(22.9%) | 55(35.0%) |
| My experience | 4(2.5%) | 1(0.6%) | 5(3.2%) |
| *Information aiding decision making* |  |  |  |
| Nutritional Information | 2(1.2%) | 3(1.8%) | 5(3.0%) |
| Lab test results | 9(5.5%) | 2(1.2%) | 11(6.7%) |
| Blood pressure check-ups results/symptoms | 5(3.0%) | 2(1.2%) | 7(4.3%) |
| Immunization information | 6(3.7%) | 4(2.4%) | 10(6.1%) |
| Information on Danger Signs | 9(5.5%) | 3(1.8%) | 12(7.3%) |
| Info. on types of services offered | 7(4.3%) | 3(1.8%) | 10(6.1%) |
| All the above | 4(2.4%) | 9(5.5%) | 13(7.9%) |
| Not sure | 31(18.9%) | 47(28.7%) | 78(47.6%) |
| *Decision making autonomy* |  |  |  |
| Yes | 41(25.2%) | 25(15.3%) | 66(40.5%) |
| No | 41(25.2%) | 56(34.4%) | 97(59.5%) |
| *Who makes your decisions* |  |  |  |
| Spouse | 12(14.6%) | 19(23.2%) | 31(37.8%) |
| Midwife | 15(18.3%) | 3(3.7%) | 18(22%) |
| Peer mothers | 2(2.4%) | 5(6.1%) | 7(8.5%) |
| Village health teams | 13(15.9%) | 13(15.9%) | 26(31.7%) |
| *Why don't you make ANC decisions* |  |  |  |
| I do not know much about ANC | 16(16.5%) | 32(33%) | 48(49.5%) |
| My husband provides the money | 14(14.4%) | 17(17.5%) | 31(32%) |
| I do not know | 7(7.2%) | 5(5.2%) | 12(12.4%) |
| I cannot make good decisions | 2(2.1%) | 1(1%) | 3(3.1%) |
| It's my culture | 2(2.1%) | 1(1%) | 3(3.1%) |
| *Challenges prohibiting ANC access* |  |  |  |
| Lack of information about ANC services | 36(22%) | 31(18.9%) | 67(40.9%) |
| Lack of money | 13(7.9%) | 11(6.7%) | 24(14.6%) |
| Long distances | 5(3%) | 16(9.8%) | 21(12.8%) |
| Inadequate and poor services at hospitals | 26(15.9%) | 17(10.4%) | 43(26.2%) |
| Cultural Inclination | 2(1.2%) | 6(3.77%) | 8(4.9%) |
| None of the above | 1(0.6%) | 0(0%) | 1(0.6%) |
| *Previous ANC attendance* |  |  |  |
| Yes | 59(37.8%) | 29(18.6%) | 88(56.4%) |
| No | 23(14.7%) | 45(28.8%) | 68(43.6%) |
| *Mothers with mobile phones* |  |  |  |
| Yes | 62(38%) | 67(41.1%) | 129(79.1%) |
| No | 21(12.9%) | 13(8%) | 34(20.9%) |
| *Access to ANC services using any ICT's* |  |  |  |
| Yes | 9(5.5%) | 6(3.7%) | 15(9.2%) |
| No | 74(45.4%) | 74(45.4%) | 148(90.8%) |

**Figure 1:**
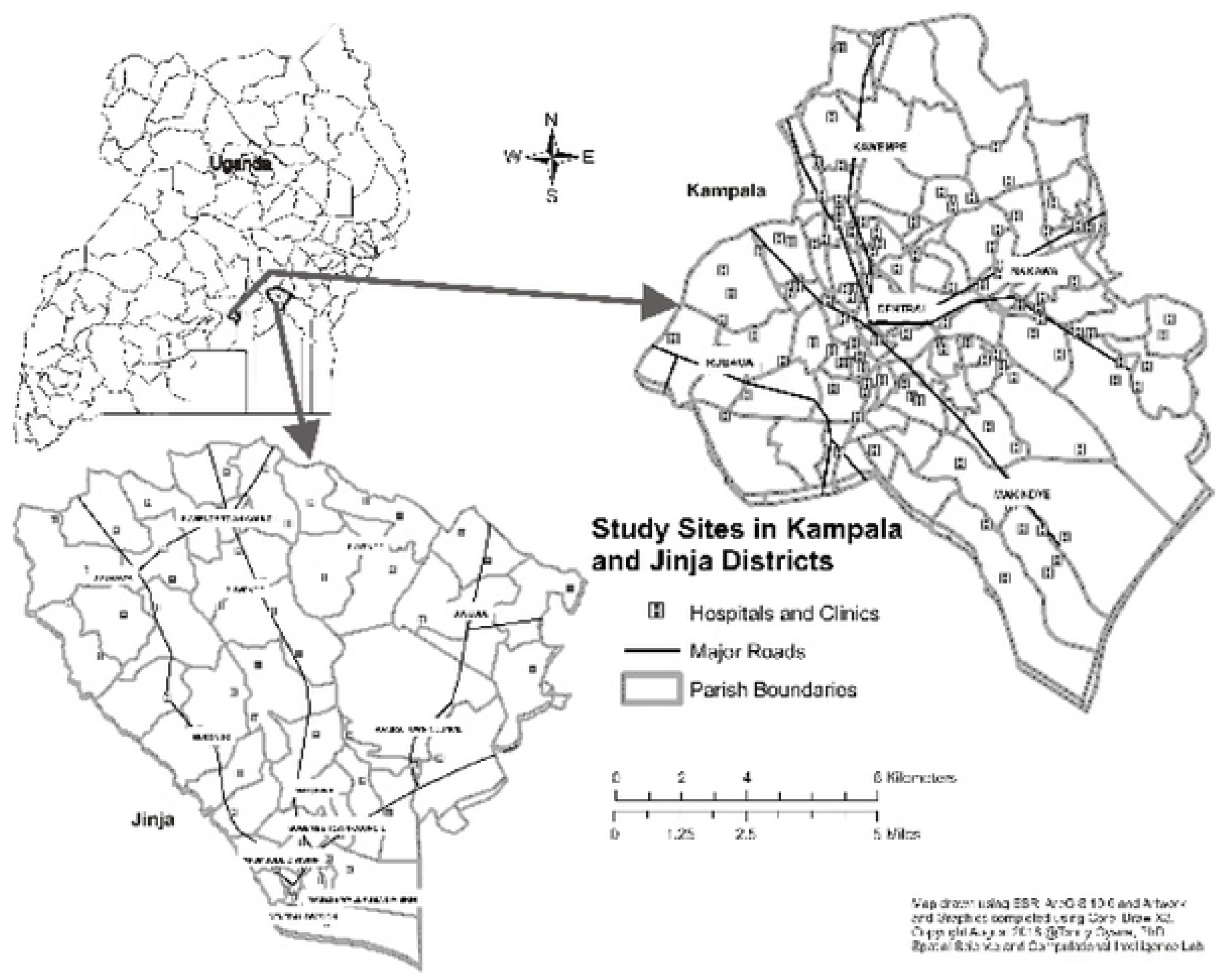
Study area map showing locations of study sites, healthcare centers and access.

**Figure 2:**
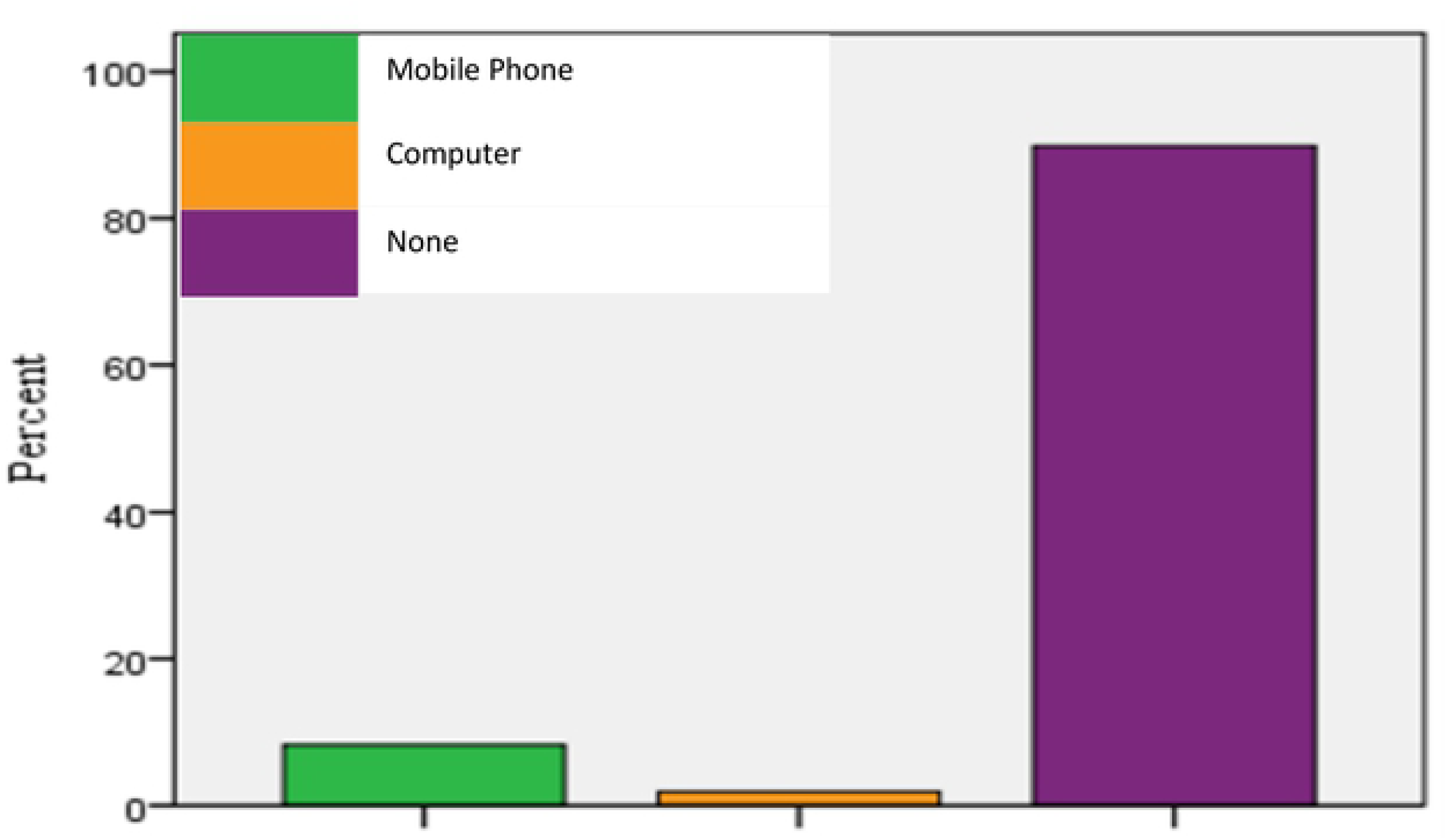
Technologies used to access ANC

**Figure 3:**
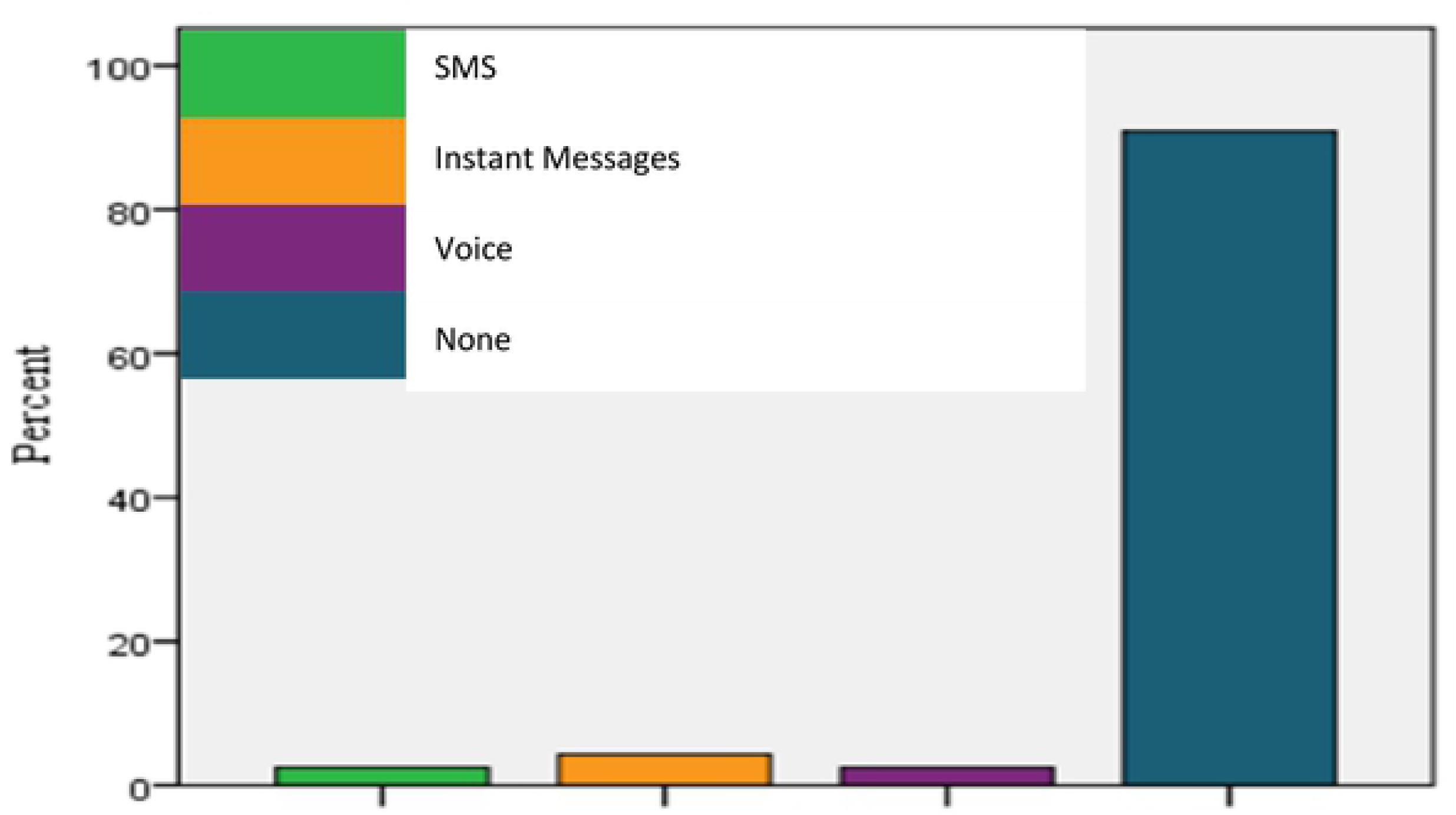
ICT services aiding ANC

**Figure 4:**
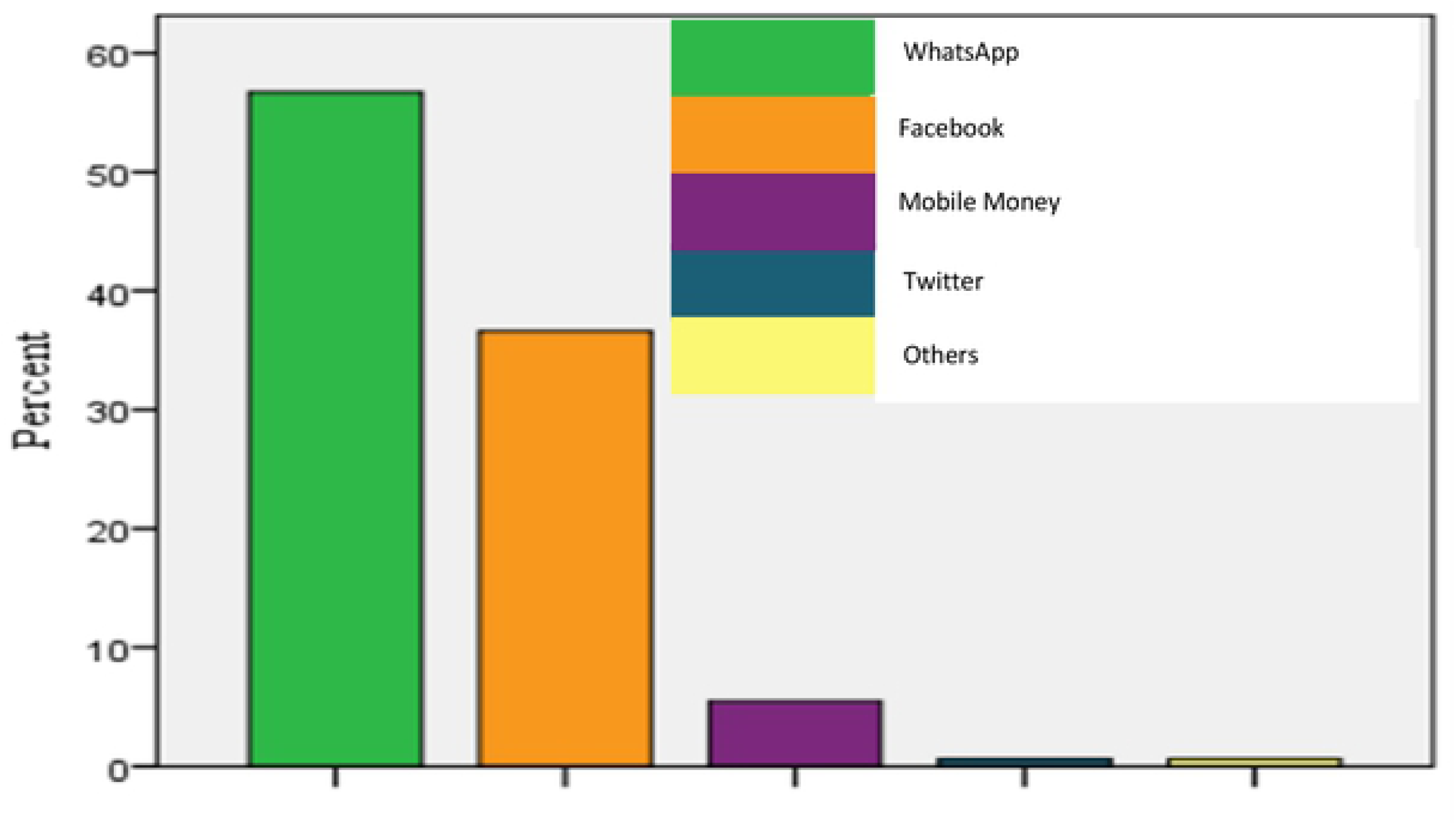
Other uses of mobile phones

**Figure 5:**
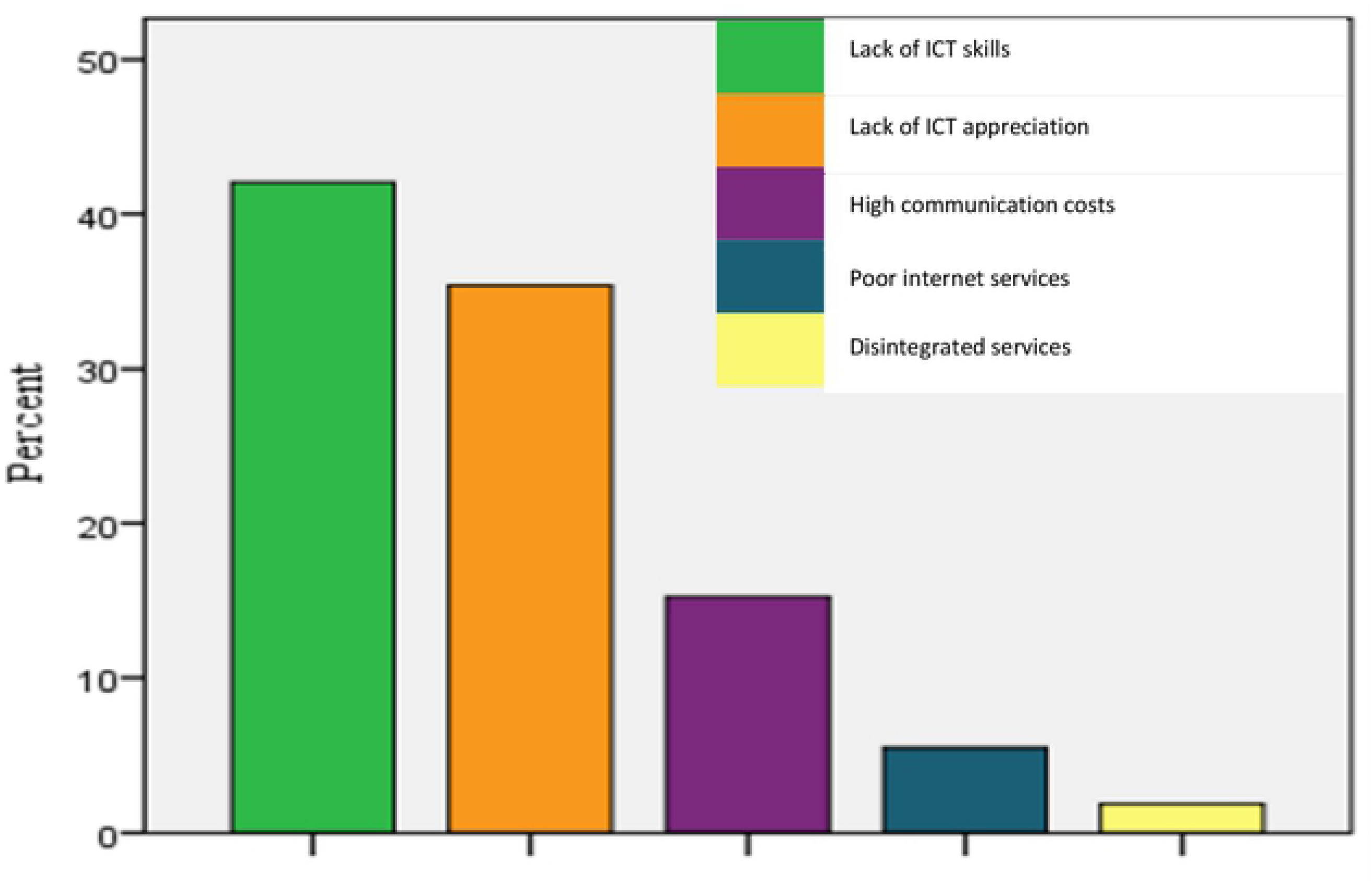
Barriers limiting uptake of ICT

### Qualitative Data

Ten women were randomly selected from the 164 and engaged further in a focus group discussion (FGD) for approximately 80 minutes. Purpose was to ascertain the nature and types of decisions that mothers routinely engage in while pregnant. The question that guided the discussion was: 1) what decisions do you usually make while pregnant? Mothers could not straight away point out the different types of ANC decisions. However, basing on the different opinions, and after a word by word thorough scrutiny and deductions from their responses, the six categories of decisions summarized below were arrived at: facility-based decisions, care decisions, nutrition, birthing, attendant-based and lifestyle decisions.

Responses from mothers are ranked according to the frequency a particular response was mentioned during the FGD. This is followed by an explicit explanation of the mothers’ opinions and experience. Excerpts from the FGD and anecdotal evidence were integrated in the narrative to give a reader a clear perspective of the mothers’ experience. Below is a summary of responses from mothers followed by an elaborative narrative of their experience.

When asked about the types of decisions they regularly engage in while pregnant, all participants agreed to have at least engaged in two of the decisions summarized in the list above. Except for the six, the rest acknowledged participating in *facility-based decisions* which entailed *“where to go for ANC, emergency care, delivery, newborn care or postnatal care”.* One participant offered a statement that echoed the sentiments of others; this statement sums up the ease with which some mothers made facility-based decisions.

> “Every time I wanted to go to hospital for ANC, I just informed my husband and he gave me money for transport”.

Single mothers made their choice of hospitals as one mother narrated *“…I am a single mother, so I decide which hospital to go to …”.* However, the choice of a health facility was generally influenced by the *“distance”, “cost of services”* and *“the type of services offered”* as narrated by some mothers.

Other mothers cited participating in *care decisions* which, to our understanding, meant *ANC decisions* and *emergency obstetric care (EMOC) decisions.* More than half of the mothers agreed to not participating in these decisions. One mother particularly acknowledged being in charge of *“where, when and what ANC services they sought for”.* Almost all (9 out of 10) mothers agreed not to participate in EmOC decisions stating that *“…they sometimes did not know what constituted an emergency…”* others had *“…domineering spouses….”* while others “…..*thought it was normal to experience certain conditions while pregnant…”* so these factors delayed the mothers’ decision to seek care.

When engaged further, more than half (7 out of 10) of the mothers acknowledged deciding *what to eat, when to eat, how and the quantity*. We collectively termed these as *nutrition decisions*. The rest acknowledged not deciding their meals, and one mother in particular described this as a *“husband’s right”,* in her narration she stated *“the husband is the head of the family and he provides the money to buy the food, so he has full autonomy over what we eat*.

Contrary to this statement, one mother in her statement that supported the opinions of others, noted that *“….it’s my role as a home maker to decide the best meal to have because my condition (pregnancy) dictates what meal we shall have in a given day….”*

At the time of delivery, many mothers (8 out 10) acknowledged deciding on how they would deliver. We referred to these decisions as *“birthing decisions”.* Of the eight mothers, 6 opted for normal delivery

> In my third trimester, I start using local herbs (commonly known as emmumbwa) to soften my pelvic bones because I cannot accept to be cut (referring to a C-section). I try as much as possible during labour to have the baby normally.

while the 2 went with caesarean section. A C-Section delivery was not an option because of the cost implication such that many mothers opted for normal delivery even when it was eminent that normal delivery wasn’t possible. The opinions of the different participants are captured in the comments of one of the participants who narrates:

Expectant mothers consider normal birth delivery as an *act of valor*. Some mothers especially from urban and affluent families to opted for SBA. We referred to these decisions as *attendant-based decisions*. Except the 6, 4 mothers had their personal gynecologists. This was attributed to the fact that these mothers did not want to go to general hospitals because in their opinion, *“…. they were too congested…” or “…care providers were insensitive and rude…”, or “.services were poor…”.* One of the mothers, in her opinion emphasized *“the lack of empathy with most care providers in these general hospital…”* and thus chose a private care provider even if it meant *“spending a lot more than usual”.* Mothers that could not afford a private gynecologist, went to general hospitals and had no choice of who was to assist them during delivery. One mother in particular stated *“…whoever I find in hospital is the one who helps me in delivery…”* and another mother supplemented *“…I have no choice on a midwife to assist me because one is lucky to find one who is willing to help you…”*

Participants indicated that they engaged in exercising, yoga, swimming, walking while pregnant. Those who drank alcohol and smoked prior to conception stopped when they conceived while others (2 out of 5) indicated taking folic acid before conception. In this study, we referred to these as *lifestyle decisions*. A few mothers pointed out their urge to stay healthy for the sake of their unborn babies. While one participant narrated that *“yoga relaxes her”* another mother admitted to *“…quitting smoking the moment she discovered she was pregnant…”*

The common threads from the qualitative data collected from expectant mothers is summarized below:

- Key factors that determine their choice of health care center are distance, cost and service type.
- Barriers to ANC decisions are spouses and inadequate knowledge on what constitutes an emergency.
- Nutrition decisions were positively reviewed in their favor due to direct home involvement in meal preparations.
- Normal delivery was the preferred method of delivery.
- SBA was mainly determined by the care providers, especially by mothers with prior experience in a general hospital. A small number of mothers who could afford a private specialist chose gynecologists as their preferred SBA.
- Staying healthy and fit was important for a few mothers and their unborn babies.

## Discussion

This study has four major findings: 1) the lack autonomy to influence ANC decisions for majority of mothers, 2) the general lack of ANC information, 3) the fundamental role of midwives and VHT’s as major sources of ANC information, and 4) the limited use of ICT’s especially mobile phones to aid decision making.

Our findings are consistent with previous studies [16, 19, 20, 21, 22, 23, 24, 25, 26, 27, 28, 29, 30, 31, 32, 33]. This study found a strong association between ANC decision-making autonomy and women with higher education (χ^2^ = 8.63, *ρ* = 0.035), married (χ^2^ = 4.1, *ρ* = 0.043) and mature (36–50) (χ^2^ = 8.81, *ρ* = 0.032); educated women were more likely to influence their decisions compared to their less educated counterparts. On average, majority of the respondents in both regions had less decision-making autonomy especially women from Jinja. Generally, spouses, midwives and VHT’s greatly influenced mother’s ANC decisions. This supports previous studies in Ghana and Bangladesh that noted husbands, mothers-in-law, family and community members being principal decision makers especially in cases of EmOC [21, 23]. This over-dependence on VHT’s for ANC decisions among less educated mothers is a barrier to ANC access. Lack of knowledge and information was the major hindrance to participation to decision making, which was also emphasized in the WHO report as one of the reasons why women in the Sub-Saharan region did not access ANC services [32]. Spouses dictated when and where to do routine ANC, where to go in case of an emergency, delivery and sometimes influenced the diet of their wives because they provided therefore exercising power, as a house head was unquestionable [33]. Also, care providers largely influenced how women were to give birth.

Suffice to say, the lack of decision-making autonomy to seek care was seen in almost half of the respondents. This can be corroborated in other similar studies [23, 31]. The lack of autonomy coupled by the lack of information, money, poor services and long distances to health facilities culminated to the non-attendance of ANC. These inhibiting factors were also common in a study conducted in Dokolo, Rwashamaire, Kayunga and Budadiri districts in Uganda [24]. During pregnancy, mothers interact with different people like VHT’s, midwives, spouses, peer mothers among others. This interaction indicates that mothers don’t make decisions in solitude. However, mother’s decision-making autonomy should be enabled, not crippled. An interesting finding which has not been reported elsewhere was the different types of decisions mothers routinely engage in which include; facility-based, nutrition, lifestyle, birthing, attendant and care-based decisions.

Expectant mothers relied more on midwives and VHT’s for information and other maternal health related services. This finding is consistent with another study [21]. However, in our study, expectant mothers in Kampala relied heavily on midwives for ANC information while those in Jinja depended more on VHT’s. This can be corroborated by previous studies conducted in Kamuli, Pallisa and Kibuku districts where the major source of information for maternal and neonatal health were VHTs [19]. Peer mothers also played a fundamental role of providing ANC information to expectant mothers especially in Jinja.

Fifty percent of expectant mothers who participated in this study acknowledged some information that was relevant in their routine ANC decision making practices. However, more than two-fifth demonstrated a clear lack of knowledge on ANC issues, which was consistent with previous studies [22, 30]. This shows that mothers are ignorant of their ANC information needs. This was more evident among women from Jinja. The general lack of knowledge is consistent with other studies conducted in Busia, Gulu and South-Western [16, 25, 28].

Notably, findings indicated that majority of the women were not using ICT’s in their routine ANC decision making practices even though many had mobile phones. A few of the mothers that managed to use ICT’s for ANC relied on mobile phones. Apart from voice services, mothers largely used their mobile phones for social media services like WhatsApp and Facebook for social interactions but not maternal issues. This relates to a similar study conducted in Ghana [27]. Although the adoption of ICT services in enhancing access and use of ANC services was still very low, which is consistent in other studies [25], a few mothers used voice, SMS and instant messages in their daily ANC routine practices with either midwives or VHT’s. The major reasons that inhibited mothers’ use of ICT’s in their routine ANC practices included the lack of technical skill to use the technologies, the limited knowledge of the capabilities of ICT’s and the high data costs, which is consistent with other studies [20, 29].

### Implications

There are two implications for this study. Firstly, the need to use the data and knowledge developed to enhance ICT use in ANC decision making practices among expectant mothers in Uganda. The decision to utilize ANC services varied greatly along social-economic and demographic factors. It was found out that expectant mothers especially those from rural settings were less aware of ANC information needs and largely depended on their spouses, VHT’s and peers for information and decision making.

Secondly, to create favorable policies that encourage expectant mothers to seek early ANC, especially in cases of emergency, to reduce the risk of maternal and/or neonatal deaths.

Findings suggest that the lack of information and confidence among mothers, husband’s financial support and cultural factors largely contributed to the low decision-making autonomy. This should be mitigated through health education awareness programs.

### Recommendation and Future Direction

To improve on maternal health outcomes, there’s a need to empower expectant mothers to make decisions to seek care from skilled birth attendants. Additional research is required to determine the effectiveness of ICT services to support ANC with goal of closing the gap between the service providers and consumers. Our findings show a strong link between expectant mothers and midwives/VHT. Interventions to strengthen the interaction and synergy between mothers and these stakeholders to enable seamless exchange of information and collaboration may create an avenue for mothers to have a certain degree of autonomy over their decision-making practices.

## Acknowledgment

We would like to extend our sincere gratitude to the staff, management and colleagues of Kampala City Council Authority and Makerere University College of Computing and Information Sciences, Kampala, Uganda. Special thanks to our research assistants, respondents and community health workers who provided critical data and support through their participation

